# Conserved transcriptomic profile between mouse and human colitis allows temporal dynamic visualization of IBD-risk genes and unsupervised patient stratification

**DOI:** 10.1101/520379

**Authors:** Paulo Czarnewski, Sara M. Parigi, Chiara Sorini, Oscar E. Diaz, Srustidhar Das, Nicola Gagliani, Eduardo J. Villablanca

## Abstract

Despite the fact that ulcerative colitis (UC) patients show heterogeneous clinical manifestation and diverse response to biological therapies, all UC patients are classified as one group. Therefore, there is a lack of tailored therapies. In order to design these, an unsupervised molecular re-classification of UC patients is evoked. Classical clustering approaches based on tissue transcriptomic data were not able to classify UC patients into subgroups, likely due to associated covariates. In addition, while genome wide association studies (GWAS) have identified potential new target genes, their temporal dynamic revealing the optimal therapeutic window of time remains to be elucidated. To overcome the limitations, we generated time-series transcriptome data from a mouse model of colitis, which was then cross-compared with human datasets. This allowed us to visualize IBD-risk gene expression kinetics and reveal that the expression of the majority of IBD-risk genes peak during the inflammatory phase, and not the recovery phase. Moreover, by restricting the analysis to the most differentially expressed genes shared between mouse and human, we were able to cluster UC patients into two subgroups, termed UC1 and UC2. We found that UC1 patients expressed higher copy of genes involved in neutrophil recruitment, activation and degranulation compared to UC2. Of note, we found that over 87% of UC1 patients failed to respond to two of the most widely-used biological therapies for UC.

This study serves as a proof of concept that cross-species comparison of gene expression profiles enables the temporal annotation of disease-associated gene expression and the stratification of patients as of yet considered molecularly undistinguishable.

## Introduction

Ulcerative colitis (UC) is a type of inflammatory bowel diseases (IBD) that is mostly restricted to the colon and is characterized by changes in the mucosal architecture, epithelial function, increase in immune cell infiltration and an elevated concentration of inflammatory cytokines. Symptoms include diarrhea, abdominal pain, rectal bleeding, lack of appetite and fatigue, all of which significantly affect patient’s quality of life. UC is recognized as a heterogeneous disease, presenting diverse macroscopic features, symptoms, grads of inflammation and colonic affected areas ^1,2^.

Although there is no definitive cure for UC, there are biological therapies available which target the inflammatory response during UC by means of inhibiting pro-inflammatory cytokines or by blocking immune cell migration ^3^. Among these, the most frequently used biological therapies in UC patients block tumor necrosis factor (TNF) with anti-TNF antibodies (such as infliximab, IFX) ^4^ or leukocyte migration (such as vedolizumab, VDZ) ^5 6^. However, about 35% ^4,6^ and 50% ^5,6^ of patients poorly achieve clinical response to IFX and VDZ, respectively. Patients that do not respond develop adverse effects, most notably increased risk of infections, thus requiring continuous medical monitoring and ultimate surgical intervention ^7,8^.

In an attempt to identify genes/pathways as a potential novel therapeutic target, genome wide association studies (GWAS) have identified more than 200 polymorphisms associated with higher susceptibility to IBD ^9,10^. However, the function and temporal expression of IBD risk genes during experimental colitis are yet to be elucidated ^9,10^.

Furthermore, while there is an obvious clinical heterogeneity among UC patients as seen for example by the location affected (i.e. distal colitis, left-sided and pancolitis, and responder and non-responder) and the extent of the severity, initial treatment for these patient subgroups is identical and modified only if the patients have not responded ^6,8^. Biomarkers that could distinguish the different entities of the UC spectrum are currently lacking and they are required in order to achieve the highly needed stratification of UC patients into molecularly functional subgroups ^8,11^. Moreover, an unbiased stratification of UC subtypes has never been accomplished at the molecular and functional levels. Here, using transcriptomic data from a well-characterized experimental model of colitis we were able to identify conserved genes between mouse and UC patients. As a result, we were able to gain insights into IBD-risk gene kinetics and to molecularly stratify UC patients in an unsupervised manner.

## Results

### Human UC is highly variable at the transcriptome level

In order to molecularly stratify UC patients into subgroups, we combined 4 publicly available human UC cohort datasets (n=102 patients), in which transcriptomic microarrays of total colonic biopsies was performed ^12–15^ (**Table 1** and **Fig S1**). We ranked genes using the top 100 most variable genes and further tested whether molecular subgroups exist (**Fig 1a**). Analysis by visual assessment of cluster tendency (VAT)^16^ indicated that biopsies presented high inter-sample dissimilarities (**Fig1b**), suggesting a poor overall tendency to form consistent clusters. Dimensionality reduction analysis by tSNE using the top highly variable genes also indicates the formation of a single group with no apparent subdivisions (**Fig 1c**). Then, we further statistically tested whether multi-cluster substructures were present in the dataset, since most clustering algorithms define subgroups even on random noise ^17–19^. However, bootstrapping analysis using the Hartigan’s Dip test ^19,20^ presented a low cluster substructure trend (p > 0.9), regardless of the gene ranking metrics used (**Fig 1d**). Independently of the clustering tendency results, we forced patient subdivision using hierarchical clustering and tested for cluster stability using bootstrapping ^17,18,21^. In line with previous results, formed clusters were highly unstable using the list of highly variable genes (AU ≈ 0%) (**Fig 1e**). These results indicate that without prior knowledge of patient subdivision, standard gene ranking strategies do not allow clustering of UC patients into molecularly distinct subgroups.

**Figure 1.**
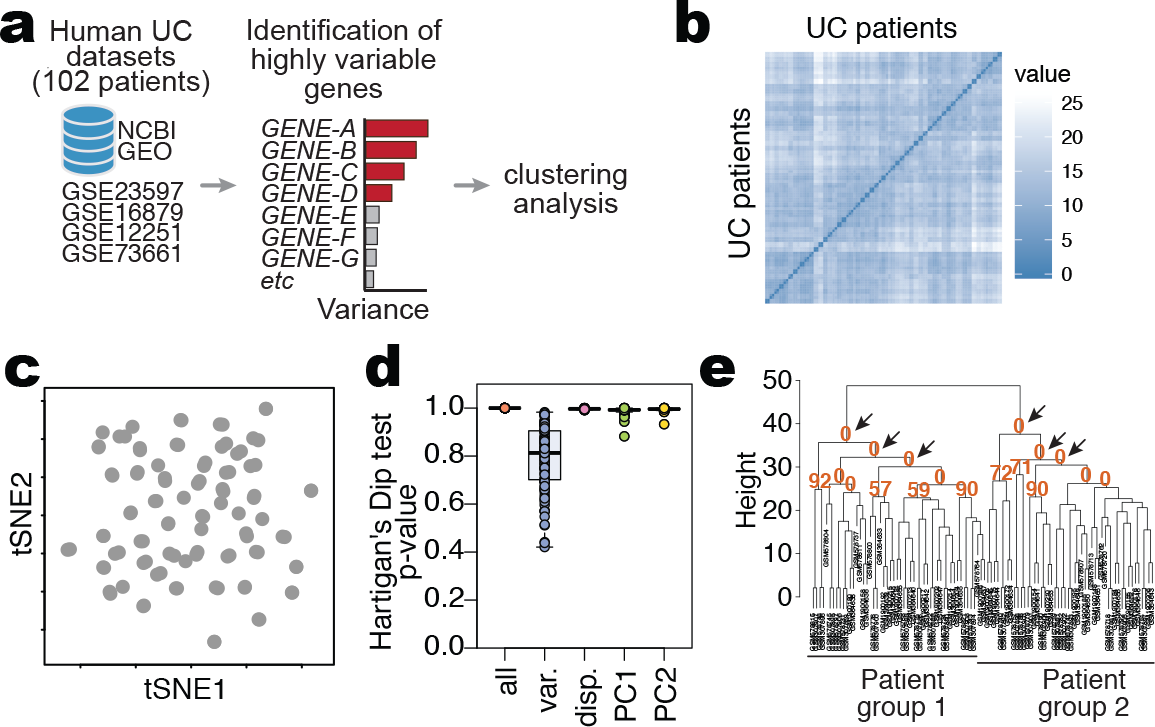
Human ulcerative colitis transcriptional profiles cluster patients into molecular subgroups. (**a**) Schematic representation of the strategy used for patient group identification, in which four publicly available datasets were combined. Gene ranking was done using the most variable genes in the human dataset, which were used for clustering analysis. (**b**) Sample dissimilarity heatmaps for visual analysis of clustering tendency (VAT), comparing the human dataset using the top 100 variable genes. (**c**) tSNE plot using the top 100 variable genes in the human dataset. Each point represents a patient sample. (**d**) Hartigan’s Dip test for clustering tendency using all genes in the dataset, the top 100 variable genes, the top 100 highly dispersed genes or the top 100 leading genes in the principal components. (**e**) Bootstrapping analysis of hierarchical clustering, comparing the human dataset using the top 100 variable genes in the human dataset. Numbers in orange indicate the approximately unbiased (AU) p-value, shown as a percentage. AU closer to zero indicates a cluster with low stability.

### Time-series reveals processes underlying colon inflammation and repair

One cause of such inter-patient variability can be attributed to the sampling procedure, which contributes largely to the total data variance and masks real biological differences ^22,23^. To overcome the total data variance, we sought to identify the genes that contribute to inflammation in an independent and unsupervised manner. To this end, we focused the analysis on a list of evolutionarily conserved genes that best discriminate the nuances of inflammation in a well-characterized colitis mouse model ^24^.

To identify these evolutionarily conserved genes, we first elucidated through an unbiased manner which genes and pathways are differentially regulated during mouse colonic inflammation followed by a tissue regeneration phase. In particular, we took advantage of the widely used dextran sodium sulfate (DSS)-induced model of colitis. This model is one of the few characterized by a phase of damage followed by a phase of regeneration. Therefore, this model gave the possibility to identify also sets of genes essential in the regeneration phase, a key step towards the resolution of the inflammation. In short, mice were exposed to DSS in the drinking water for 7 days, then allowed to recover for the following 7 days. During this period, we collected colonic tissue samples every second day to then be analyzed by RNA sequencing (RNA-seq), histology and flow cytometry (**Fig 2a and Fig S2**). First, we confirmed that 7 days of DSS exposure resulted in continuous body weight loss and acute disease severity until day 10 to then initiate the recovery phase (**Fig S2a-b**). Histological analysis confirmed epithelial damage, such as desquamation of the epithelial layer on day 6 (**Fig S2c**), while labeling proliferating cells within crypts (Ki67 staining) indicated disrupted crypt architecture by day 6 and restoration by d14 (**Fig S2c**). Loss of the epithelial cells 117 (CD45^neg^EpCAM^+^) by day 7-10 and restoration by day 14 was further confirmed by flow cytometry (**Fig S2d**). To test whether the epithelial barrier integrity was restored by day 14, we gavaged FITC-dextran and measured its concentration in the serum. We detected higher FITC-dextran concentrations on day 7, which indicates barrier disruption, whereas basal levels were detected by day 14 indicating restoration of the barrier integrity (**Fig S2e**). Thus, on the basis of this characterization we will refer to d6-d10 and d12-d14 as acute phase and recovery phase, respectively.

**Figure 2.**
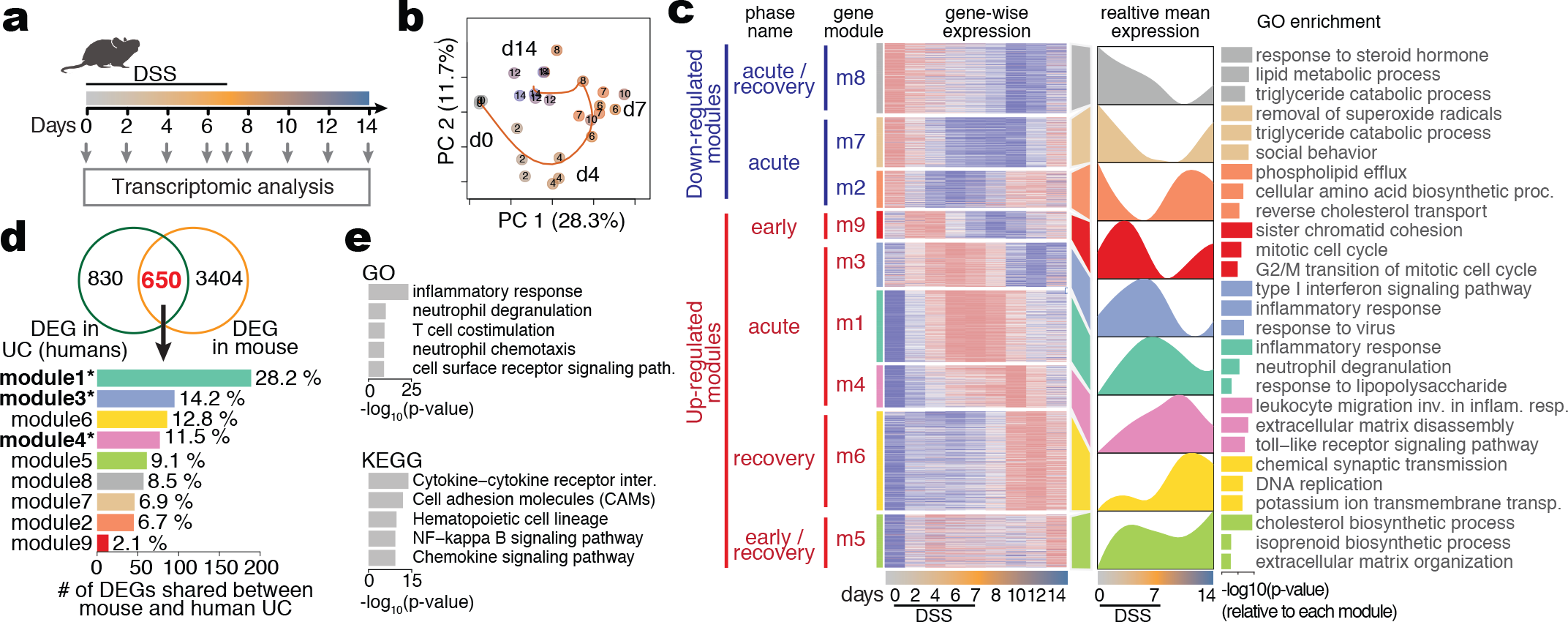
Unbiased characterization of the DSS colitis reveals conserved inflammatory signature between mice and humans. (**a**) Schematic illustration of the experimental design. Mice received DSS in drinking water for 7 days, after which the treatment was replaced with water. Samples were collected at indicated time points. (**b**) PCA on differentially expressed gene counts. Samples were color-coded according to their respective day of collection (from grey to orange to blue). The percentage of variance explained by the respective principal component indicated in parenthesis. (**c**) Clustered heatmap of all differentially expressed genes (left). The mean expression of each gene module is shown (right). Functional annotation of genes in each cluster was done based on Gene Ontology (GO) enrichment. Only the top 3 enriched processes are shown, sorted by P-value. (**d**) Venn diagram comparing the list of DEGs in treatment-naive UC and the DEGs identified in mouse DSS colitis (upper). Among the 650 genes shared among those lists (in red), the number and percentage of genes found in each module identified in our mouse dataset (lower). Modules highlighted in bold are the ones enriched for inflammatory terms in **Fig 2c**. (**e**) GO and KEGG enrichment analysis out of the 650 shared genes identified in (**d**), sorted by P-value.

Next, we performed a RNA-seq analysis from colonic samples throughout the experiment and computed differentially expressed genes (DEGs) taking the complete kinetics of expression into consideration for p-value estimation using EdgeR ^25^ (see Methods). A detailed list of all genes found differentially expressed is available for further exploration (**Table S1**). Principal component analysis (PCA) on DEGs revealed that samples displayed a sequential temporal path in PCA space, starting on day 0, passing through day 7 (acute) and ultimately reaching day 14 (recovery) (**Fig 2b**). Of note, samples from day 14 did not reach the same gene expression profile compared to day 0, suggesting that complete molecular restoration was not reached by day 14. We observed that over 70% of the variance among the differentially expressed transcripts is retained in the first 5 principal components (PCs) (**Fig S3a**), and that each principal component corresponds to a unique expression kinetic through the time course of DSS-colitis (**Fig S3b**). For instance, the variance explained by PC1 peaked at the acute phase and returned to almost normal levels on day 14 (recovery), capturing most of the variance related to inflammatory genes that peaked from days 7 to 10, such as *Ly6g*, *Reg3b*, *Reg3g*, *S100a8*, *S100a9*, *Mmp3*, *Mmp8*, *Mmp10* (**Fig S3b and c**). On the other hand, the variance explained by PC2 peaked on day 4 during DSS administration, to return close to normal by day 7, thus, capturing most of the variance related to genes expressed during initiation of inflammation, such as *Mcpt1*, *Mcpt2*, *Mmp3*, *Mmp10*, *Il11*, *Scnn1g* and *Best2* (**Fig S3b and c**). These results indicate that several of the genes modulated between days 4-10 are related to inflammation and together contribute the most to the variance in the dataset.

By using hierarchical clustering on the spline smoothed gene expression of DEGs, we were able to classify the gene expression into 9 modules (**Fig 2c**). For further exploration, expression values for all genes in each module are available (**Table S1)**. Three gene modules (m2, m7 and m8) were down regulated during the acute and recovery phases of DSS-induced inflammation, with lowest peak on days 6, 10 and 12, respectively. GO and KEGG enrichment analysis suggest that these modules represent genes mainly involved with epithelial cell functions, such as PPAR signaling (*Acsl1*, *Fabp1*), small molecule metabolism (*Sult1a1*, *Sult1b1*) and fat digestion and absorption (*Paqr8*, *Clps*, *Pla2g3*) (**Fig 2c and Fig S4a**).

On the other hand, six modules (m9, m3, m1, m4, m6 and m5) were up-regulated over the early, acute and recovery phases of DSS-induced inflammation, peaking on days 2, 6, 7, 10, 12 and 14, respectively. Among those, processes such as cytokine signaling (*Il11*, *Il12b*, *Il6*, *Il1b*), leukocyte migration (*Sell*, *Ccr1*, *Ccr2*, *Cxcl2*, *Cxcr3*), neutrophil degranulation (*Ly6g*, *Itgam*, *Itgax*, *Cd300a*), matrix remodeling (*Mmp3*, *Mmp7*, *Mmp10*), response to lipopolysaccharide (*Saa3*, *Nox2*) as well as several inflammatory signaling pathways (*Stat3*, *Jak3*, *Nfkbia*, *Smad4*, *Birc3*) were enriched, suggesting the interplay of several immune cells and pathways as a cause/trigger of inflammation, especially during the acute phase (**Fig 2c and Fig S4b**). Moreover, modules m9 and m5 presented two degrees of bimodal expression pattern, peaking at day 2-4 (early phase), with slight down-regulation between days 7-10 and a second peak on days 12-14 (recovery phase). Genes in those modules were associated mainly with cell cycle (*Ttk*, *Cdc7*, *Cdc20*, *Cdc25c*, *Ccna2*, *Ccnb1*, *Ccnb2*) and cholesterol biosynthetic pathways (*Acat2*, *Sqle*, *Mvd*, *Hmgcs1*), respectively (**Fig 2c and Fig S4b**). Many other genes and GO/KEGG pathways not shown here are fully accessible for exploration of individual genes and their clusters (**Table S1, S2 and S3**). Taken together, time-series transcriptomic characterization of mouse colonic inflammation identifies distinct gene expression kinetics associated with epithelial and immune cell related pathways during the course of colitis.

### Inflammatory pathways are the most conserved between mouse and human colitis

Having characterized genes and pathways that are associated with intestinal inflammation and tissue repair during experimental colitis, we investigated whether such pathways are conserved in humans. To this end, we compared the list of DEGs from the mouse experimental colitis with the recently published list of DEGs found in newly diagnosed treatment-naïve ulcerative colitis patients ^26^. This is a cohort containing human RNA-seq data, where they report DEGs between UC patients versus healthy controls. We found that among the 4045 mouse DEG, 650 genes were also found among the list of DEG obtained comparing UC patients versus healthy controls (**Fig 2d and TableS4**). Out of the 650 genes shared between mouse and humans, 53.9% were identified in the inflammatory modules m1 (28.2%), m3 (14.2%) and m4 (11.5%) (**Fig 2d**). This suggests that acute inflammatory genes in m1, m3 and m4 are conserved between DSS-induced colitis and UC. GO and KEGG enrichment analysis revealed that those 650 genes were enriched for inflammatory pathways related to neutrophil degranulation and chemotaxis, as well as cytokine and inflammatory signaling pathways (**Fig 2e** and **TableS5**). These results showed that most of the genes/pathways conserved between experimental mouse colitis and human UC are associated with inflammatory responses.

### Forward translation from mouse to human UC patients allows the temporal classification of the IBD risk genes

To understand the temporal expression of the genes associated with the identified IBD polymorphisms (candidate IBD risk genes) ^9^, we checked the expression of genes associated with UC or CD identified by single variant fine-mapping resolution ^10^ into the list of DEGs from the mouse dataset. Out of the 233 reported candidate IBD risk genes, 40 genes presented very low or undetectable counts in the mouse dataset (i.e., *IL23R*, *SULT1A2*, *ERAP2*, *MUC19*), 118 were detected but did not have their expression altered through the development of inflammation (i.e., *TNFRSF14*, *ATG16L1*, *GPR35*, *TNFSF8*) and 75 were found among the DEGs in our mouse dataset (**Fig S5a** and **Table S6**). Among these, many IBD-risk genes with already known functions during mouse colitis were found (e.g. *IFNG*, *GPR65*, *ITGAL*, *CCL7*, *STAT3*, *FUT1*, *CD40*, *SULT1A1*, *MUC1, CARD9, IL12B, IRF1, CD5)*, being specifically present in gene modules related to inflammation m1, m3 and m4. Moreover, 26 genes of the 75 IBD risk genes found in our dataset are shared between UC and CD (i.e. *CARD9*, *SULT1A1*, *STAT3*, *GPR65*, *IL12B*), while 10 and 39 were restricted to UC or CD, respectively (**Fig S5b** and **Table S7**). In order to provide temporal information regarding the expression of IBD-risk genes during inflammation and repair, we utilized the mouse transcriptional landscape to map at which time point homolog IBD risk-genes were up− or down-regulated. Out of the 75 genes shared between mouse DEGs and IBD risk genes, 45 (60%) were mapped to modules m1, m3 and m4, which represent the acute phase of inflammation (**Fig S5c** and **Table S7**). Among them we found *Card9*, *Ifng*, *Il12b*, *Stat3*, *Stat4*, *Cd40*, which have been reported to exert functions during the acute phase of intestinal inflammation ^27–32^. By contrast, *Fut1*, *Sult1a1*, *Hes5* and *Tnfsf15* were mapped to modules m8, m7 and m2, which are down-regulated during acute inflammation, while *Rasip*, *Ntn5* and *Rtel1* matched with module 6 which is associated with genes that are up-regulated during the recovery phase after acute inflammation (**Fig S5c**). These data thus provided temporal information on when IBD risk genes are differentially expressed during damage and tissue repair, providing useful insights into their potential roles during inflammation and recovery.

### Key conserved inflammatory genes distinguish two human ulcerative colitis subgroups

Having identified genes that contribute to inflammatory pathways that are conserved between mice and humans, we next used those genes to assess whether UC patients can be subdivided into subgroups (**Table1**, **Fig 3a**). To this end, we selected the top 100 leading genes in PC1 and PC2 from the mouse colitis dataset and identified the respective human homologs (**Fig 3a**). We found that 57 genes were shared between mice and humans. Of these, only 17 genes were found among the 100 most variable genes of the human dataset (**Fig S6**), which might explains why patient classification using highly variable genes was not possible.

**Figure 3.**
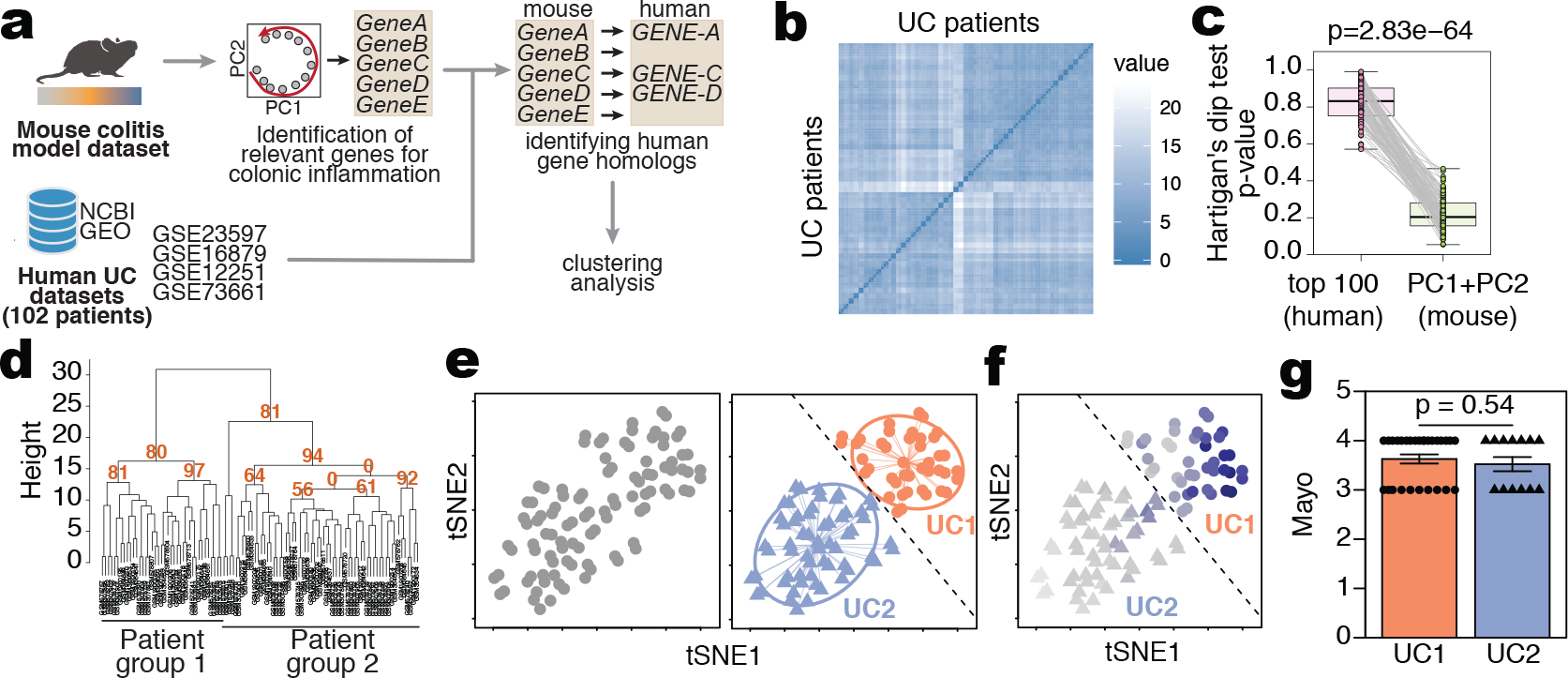
Conserved inflammatory gene signature distinguishes two UC subgroups. (**a**) Schematic representation of the strategy used for patient group identification. Four publicly available datasets were combined. Gene ranking was done using the most variable genes identified mouse dataset that had a homolog in humans. (**b**) Sample dissimilarity heatmaps for visual analysis of clustering tendency (VAT), comparing the human dataset using the top mouse gene homologs. (**c**) Hartigan’s Dip test for clustering tendency comparing the analysis using top 100 variable genes and the top mouse gene homologs. (**d**) Bootstrapping analysis of hierarchical clustering, comparing the human dataset using the top mouse gene homologs. Numbers in orange indicate the approximately unbiased (AU) p-value, shown as percentage. AU closer to zero indicates a cluster with low stability. (**e**) tSNE plot using the top variable genes identified from the mouse dataset. Each point represents a patient sample. tSNE plot showing the separation of 2 patient subgroups (left). Unsupervised hierarchical agglomerative clustering was used to automatically define patient subdivision (center). Dashed line delimits UC1 (triangle) and UC2 (circles) patients. (**f**) Average expression of mouse homolog genes used to subdivide patients (right), where dark blue colour indicates higher average expression. Dashed line delimits UC1 (triangle) and UC2 (circles) patients. (**g**) Assessment of Mayo clinical subscore in patients from UC1 and UC2. Mann-Whitney test was used for comparison.

Therefore, we performed an unsupervised analysis of the human dataset using the 57 homolog genes (**Fig 3a**). Of note, VAT analysis using these 57 homolog genes indicated the distinction into 2 major patient subgroups (**Fig 3b**), which also resulted in reduced Hartigan’s unimodality test (p < 0.001, **Fig 3c**). This indicates that by using mouse most variable genes as opposed to the sole top human variable ones, it is possible to obtain higher clustering tendency of the UC patient data. To test whether using the mouse homologs also impacted on cluster stability, we performed a bootstrapping analysis. This time, clustering using the top mouse homolog genes resulted in clusters with higher stability (AU ≈ 80%) (**Fig 3d**), compared to using the top human highly variable genes (AU ≈ 0%) (**Fig 1e**). Hierarchical agglomerative clustering using the mouse homolog genes thus defined 2 UC subtypes, namely UC1 and UC2, comprising 60 and 42 patients, respectively (**Fig 3e**). The UC1 subgroup is defined as patients presenting the higher average expression of the inflammatory genes compared to UC2 (**Fig 3f**). We also observed that neither UC1 nor UC2 subtypes were discriminated by the overall macroscopic disease severity (**Fig 3g**), suggesting that although these two UC subtypes are indistinguishable based on Mayo score, they are transcriptionally distinct.

### UC1 and UC2 are transcriptionally distinct

In order to characterize UC1 and UC2 beyond conserved genes, we performed differential expression analysis using all genes present in the human dataset. We were able to identify 205 highly differentially expressed genes, among which 187 were up-regulated in UC1 and 18 were up-regulated in UC2 (**Fig 4a**). Detailed tables with information on all DEGs comparing UC1 and UC2 are available for exploration (**Table S8 and Fig S7a**). Among those, cytokines (*TNF*, *IL11*), enzymes (*NOX1*, *MMP3*, *CYP26B1*), calcium-binding proteins (*S100A8*, *S100A9*), chemokines (*TREM1*, *CXCL8*) and other proteins related to the inflammatory response (*NR3C2*, *BCL2A1*, *PARM1*, *TNFSF13B*) were clearly able to discriminate UC1 from UC2 (**Fig 4b and Fig S7**). Enrichment analysis for cell types, GO, and KEGG pathways revealed that genes highly expressed in UC1 (187) were associated with terms related to neutrophil, neutrophil degranulation and cytokine-cytokine receptor interaction, respectively (**Fig S7b**). Venn diagram of the top enriched terms revealed many overlapping genes are shared among these pathways (**Fig 4c**), suggesting that UC1 patients present a distinct transcriptional signature enriched in neutrophil activity and cytokine signaling compared to UC2 patients.

**Figure 4.**
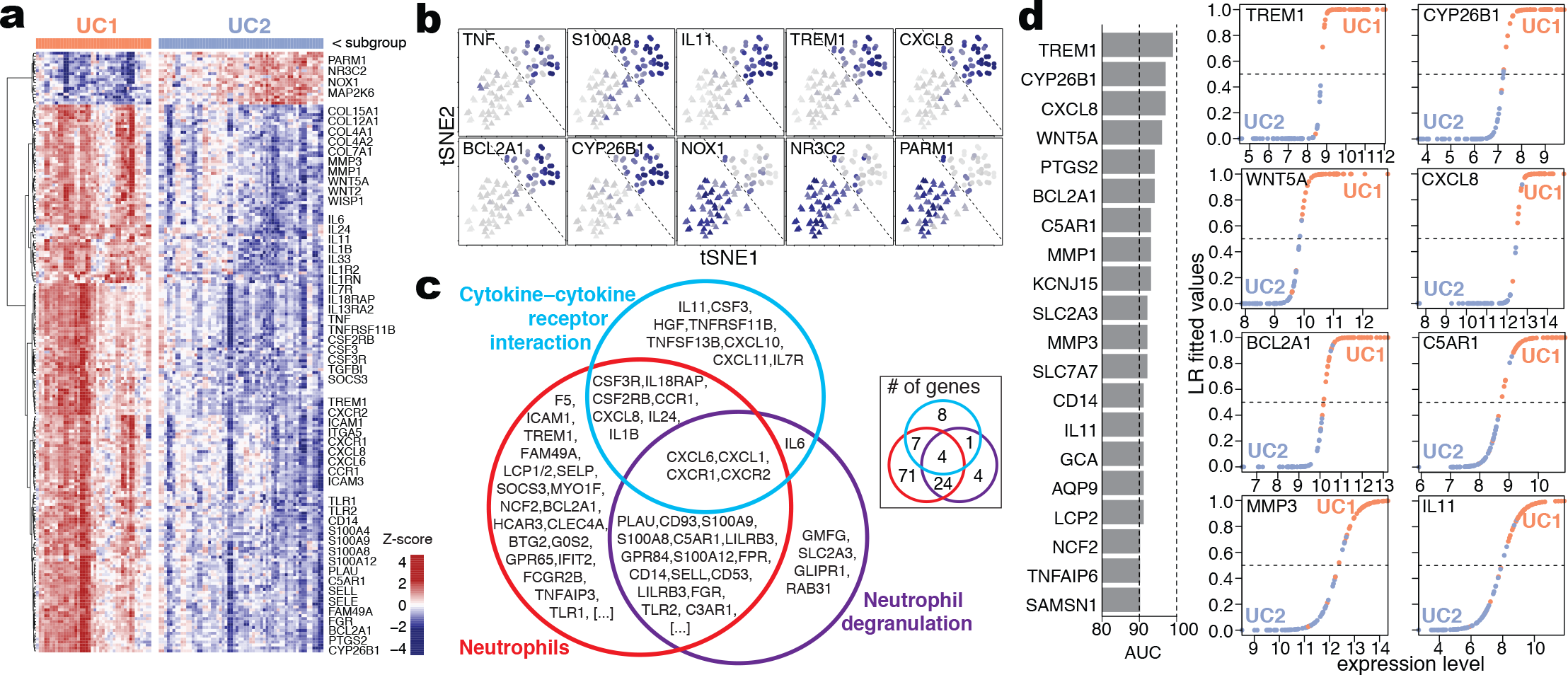
UC1 subgroup is enriched for the inflammatory signature. (**a**) Heatmap of DEGs between UC1 and UC2 patients including all genes in the human dataset. Only the selected genes are shown, grouped by functional categories and respective to the expression level. (**b**) tSNE overlay of the expression level of selected DEGs between UC1 and UC2, showing inter-patient variation. (**c**) Venn diagram of the top GO, KEGG and cell enriched terms identified from the DEGs between UC1 and UC2. (**d**) Top 20 genes ranked by area under the curve (AUC) for specificity and sensitivity to distinguish UC1 from UC2, among the list of DEGs (left). Classification was carried out using logistic regression. The fitted values of prediction are shown for selected genes (right).

We trained a logistic regression classifier using each of the DEGs between UC1 and UC2 to identify key genes that could be further used in the clinics for distinction of UC1 and UC2. Genes were tested and scored individually using the area under the curve (AUC) as a combined measure of sensitivity and specificity (**Fig 4d**). We observed that genes such as *TREM1* (AUC=99%), *CYP26B1* (AUC=97%) and *CXCL8* (AUC=97%) were among the top markers to distinguish UC1 from UC2. Other genes such as *WNT5*, *BCL2A1*, *C5AR1*, *MMP1*, *MMP3* and *IL11* also presented AUC scores above 90% and also represented good candidates for UC1 and UC2 distinction in clinical practice.

### UC1 and UC2 respond differently to biological therapies

While we stratified UC patients into two molecularly distinct subgroups, it was unclear whether UC1 and UC2 show different treatment responses to biological therapies. To address this, we used the patient-specific treatment response obtained 4 to 8 weeks after the biopsy was taken and treatment with IFX started (**Table 1**). Interestingly, we observed that on average, 70% of the patients belonging to the subtype UC2 responded to infliximab therapy (**Fig 5a**) in contrast to less than 10% of the patients classified as UC1, regardless of the dataset analyzed (**Fig 5a**).

**Table 1.** Publicly available human datasets used for classification of ulcerative colitis subtypes. Only the number of patients used for analysis are shown (inflamed mucosa before receiving any therapy).

| Dataset ID | Total | Resp | Non-resp | Ref. |
| --- | --- | --- | --- | --- |
| **Infliximab** |  |  |  |  |
| GSE12251 | 23 | 11 | 12 | 13 |
| GSE73661 | 23 | 15 | 8 | 15 |
| GSE23597 | 32 | 7 | 25 | 14 |
| GSE16879 | 24 | 16 | 8 | 12 |
| Sum | 102 | 49 | 53 |  |
| **Vedolizumab** |  |  |  |  |
| GSE73661 | 37 | 23 | 14 | 15 |

**Figure 5.**
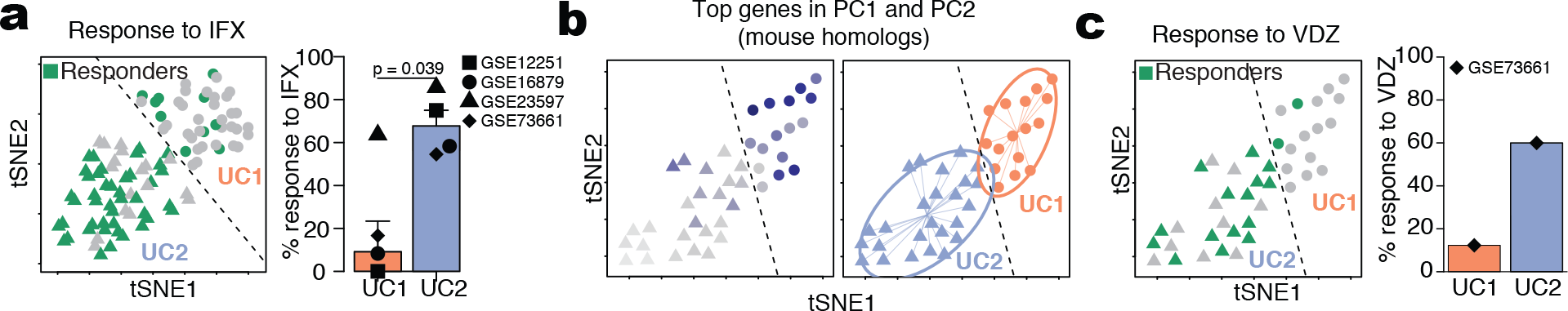
UC1 and UC2 differ on their repose to IFX and VDZ therapy. (**a**) Individual patient response to IFX therapy in each group and the percentage of patients responding to IFX in each cohort. (**b**) tSNE plot using the top variable genes identified from the mouse dataset. Average expression of mouse homolog genes used to subdivide patients (left), where dark blue colour indicates higher average expression. Unsupervised hierarchical agglomerative clustering was used to automatically define patient subdivision (right). Dashed line delimits UC1 (triangle) and UC2 (circles) patients. (**c**) Individual patient response to VDZ therapy in each group and the percentage of patients responding to VDZ in the cohort (right).

To extend the applicability of our findings, we made use of another set of UC patients receiving vedolizumab and repeated the same procedure as before (**Table 1**). Transcriptomic data from UC patients were analyzed using the most relevant genes identified in our mouse colitis model and then clustered as described above to reveal UC1 and UC2. Between them, UC1 presented a higher expression of the conserved inflammatory genes (**Fig 5b**). We observed that about 60% of the patients belonging to the UC2 subgroup responded to VDZ, in comparison to about 13% of the patients belonging to the UC1 subgroup (**Fig 5c**). Taken together, the data indicates that patients belonging to the UC2 subgroup, which present a higher percentage of response, respond to either IFX or VDZ treatment. Importantly, our approach actually allows a more accurate identification of those patients with UC1, in which 87% of the patients are refractory to both IFX and VDZ.

## Discussion

A systematic study demonstrated that biopsy sampling was the major source of inter-patient variability^22^. Therefore, such technical variations can mask real biological differences, even though UC is known to present a high level of variability in macroscopic and endoscopic scoring among patients^1,2,8^. To solve this, we limited the analysis to the relevant genes for inflammation including the phases of tissue repair and regeneration. By using the key DEGs obtained by a mouse model of colitis, we were able to “ignore” genes that were highly variable between patients (e.g. as a result of technical variation), and focus only on those that contribute to inflammation. This allowed us to temporally classify IBD risk genes and molecularly sub-classify UC patients into two subgroups; one of these characterized by genes involved in neutrophil recruitment, activation and degranulation, and by low response to biologicals.

Different experimental models to study mucosal immune processes associated with the pathogenesis of UC are available ^33,34^. Among them, the DSS-induced colitis model is broadly used due to its simplicity and applicability with different therapeutic drugs ^35^. Early studies characterized the temporal changes by qPCR for a handful of inflammatory markers ^36^, but how non-inflammatory (i.e. repair-related) genes fluctuate over time during tissue repair was unknown. Others had previously performed a kinetic microarray analysis only during the acute inflammation phase of DSS (from days 0 to 6)^37^, but whether those genes continue to be expressed during tissue repair remained unclear. Moreover, although the DSS-induced colitis model has been extensively used for the study of UC, an open reference for gene expression during intestinal inflammation and tissue repair was still missing. Here, we used a time-series transcriptional characterization of colitis, which allowed us to identify which genes contribute to most of the nuances of inflammation over time. In addition, this manuscript provides an open data source that can be further investigated by others with different questions. As an example, we provided a temporal assignment of IBD risk genes that might offer insight into their potential functions. Finally, our data show that the DSS mouse model is a relevant model for studying certain aspects of human UC.

Previous studies identified the molecular differences among responder and non-responder IBD patients ^13^. These studies were purposely biased by an *a prior*i knowledge of the responder and non-responder IBD samples. In contrast, we successfully classified the patients using a completely unsupervised approach and therefore, we have potentially identified genes that go beyond the responsiveness to the therapy by describing the molecular signature of the identified subgroups. We were able to do this by using the key DEGs found in the mouse model of colitis, by “debiasing” the human analysis, by “ignoring” genes that were highly variable between patients, and by focusing only on those genes that contribute to inflammation. Consequently, we identified two subpopulations of UC patients (UC1 and UC2).

While per definition both UC1 and UC2 subpopulations are considered inflamed, only UC1 patients present higher expression of genes associated with neutrophil degranulation and cytokine signaling, and only 10% of these patients responded to biological therapies. Similar to our results, others have shown that IL6, IL11, IL13RA, STC1 and PTGS2 were down-regulated in patients responsive to IFX ^13^ (namely UC2 in our study). Another recent report showed that the gene OSM is up-regulated in IBD patients compared to healthy controls and is predictive of anti-TNF responsiveness ^38^. However, we did not find OSM as differentially expressed between UC1 and UC2 patients. For VDZ, however, a signature for prediction of response to therapy was still missing ^15^.

The identification of UC2 which is characterized by responsiveness to both IFX and VDZ may have direct implications in the clinical setting. For example, it indicates that UC2 patients would benefit from a treatment with IFX only, since IFX therapy has a higher response rate ^6^ and is more cost-effective compared to VDZ ^39^. On the other hand, identification of non-responsiveness to both IFX and VDZ in the UC1 patient subgroup, suggests that another line of therapy should be applied. For example, we observed that the B cell activation factor (*TNFSF13B*, protein BAFF) was to be found up-regulated in UC1 patients. This suggests a potential role of B cells in UC1. Moreover, B cells are known to enhance inflammatory responses by cytokine secretion such as TNF and IL-6 ^40^, which are also up-regulated in UC1 patients. B cell depletion using anti-CD20 antibody in a small cohort showed a trend in reducing inflammation, although non-significant ^41^. However, it remains possible that B cell depletion might affect only UC1 patients, but not UC2. Similarly, we also observed that UC1 patients have a higher expression of genes involved in the JAK/STAT signaling pathway (*PTP4A3*, *SOCS3*) and cytokine signaling (*IL6* and *IL1B*), suggesting a potential role of other therapies for this subgroup, such as canakinumab (anti IL-6 mAb), siltuximab (anti IL-1β mAb), JMS-053 (PTP4A3 inhibitor) and others might apply.

In summary, we have performed an unbiased characterization of the inflammatory and tissue repair processes using a mouse colitis model, providing a useful resource for understanding colonic inflammation. Many of the genes identified in mice were also detected in human UC patients, thus allowing us to explore the temporal expression of IBD risk genes during the course of inflammation and gain useful insights into their potential function. Furthermore, they allowed us to identify for the first time two clinically relevant molecular ulcerative colitis subsets (UC1 and UC2) in an unsupervised manner. Thus, our methodology identified two molecularly distinct UC subgroups and will serve as a proof of concept for the use of transcriptomic data from highly controlled mice experiments to perform unsupervised and biologically-driven analysis of highly variable human datasets.

## Methods

All methods used in this paper are described in the Online Methods linked to this manuscript.

## Acknowledgements

We would like to thank Stefan Bonn, Samuel Huber and Charlotte Hedin for critical reading and suggestions on the manuscript. We thank Elaine Hussey for editorial assistance. EJV was supported by grants from the Swedish Research Council VR grant K2015-68X-22765-01-6, Formas grant nr. FR-2016/0005 and Wallenberg Academy Fellow (WAF) program.

## Author Contributions

PC and EJV conceived the idea and wrote the paper. PC performed bioinformatics analysis and schematic illustrations. NG and EJV provided reagents and guidance. PC, SMP, SD, CS and OED performed the experiments. PC, NG and EJV analyzed and interpreted the data. All authors contributed to manuscript writing.

## Online Methods

### 1.1. Mice and induction of DSS colitis

Female 8-12 weeks old C57BL/6J mice were obtained from ScanBur (Charles River, Germany) and housed in environmentally enriched ventilated cages under specific pathogen free conditions (SPF) at Astrid Fagræus laboratory (AFL, Karolinska Institutet) under 12h light cycle and receiving water and ration *ad libitum* (RM1(P), Special Diet Services). For induction of colitis, 2.5% w/v dextran sulfate sodium (DSS; Affymetrics) was supplemented in drinking water and given to mice for 7 consecutive days, with a change on day 3. After the treatment was ceased, mice returned to receive standard water. Mice were monitored everyday for alterations in body weight, disease activity index (DAI) ^1^. Mice were anesthetized with isofluorane and sacrificed for blood and tissue sampling. Animal experiments were done following institutional guidelines of the Stockholm Regional Ethics Committee under approved ethical permit number N89/15.

### 1.2. Mouse gene expression by mRNA sequencing

Colon samples were stored in RNAlater (Ambion) at −80°C until further use. Colonic samples were homogenized using bead-beating system (Precellys) for total RNA purification using RNAeasy kit (Qiagen) following manufacturers recommendations. RNA purity and quantity was measured by NanoDrop spectrophotometer (ThermoFisher). All samples were screened for RNA integrity check and presented RIN values above 8 on 2100 Bioanalyzer instrument (Agilent). Samples were submitted to Novogene for library preparation using TruSeq Stranded mRNA Library Prep Kit (poly-A selection) and sequencing using HiSeq-2500 platform (Illumina). Samples were sequenced using single-end 50bp sequencing^2^, aiming an coverage of 20M reads. Read quality was inspected using MultiQC^3^, trimmed with Trimmimatic^4^ and further proceeded for abundance estimation using Kallisto^5^. Further data analysis was done in R programming language (Rstudio). Genes with absolute read count less than 5 in at least 3 samples were considered with low expression and filtered out. Differences in tissue cell composition that could affect transcriptional pools were balanced by means of removing unwanted variation based on negative control genes using the RUVg function implemented in RUVseq package^6^. Analysis revealed that library sizes strongly correlated with several known intestinal housekeeping genes, such as *Hprt* (r=0.87) and *Gapdh* (r=0.85), but not *Actb* (r=0.68). Moreover, genes such as *Cd63* (0.94), *Trappc* (r=0.97), and *Cpped1* (0.97) and *Slc25a3* (r=0.96) correlated even more strongly to the library sizes, indicating potentially novel housekeeping genes during colonic inflammation. Negative controls genes were thus defined as genes with positive Pearson correlation above 0.9 to their respective sample library sizes. Estimated unwanted variation vectors were then used as covariates for calculation of differentially expressed genes (DEGs) using EdgeR package^7^.

EdgeR is specialized in performing time-series differential expression by means of generalized linear model (glm) function^8^, where time points were parsed as independent factors in the contrast matrix, thus allowing detection of differentially expressed genes at any given time point. Genes were considered differentially expressed when the overall false discovery rate (FDR) < 0.01 and at least one time-point had fold change > 1.5. DEGs identified in this manner were used for dimensionality reduction by principal component analysis (PCA), from which gene-wise contribution to the total variation can be calculated.

Identification of gene modules was done based on smoothed temporal expression curves^9^. Briefly, gene-wise log fold changes were smoothed using spline curves and further grouped into modules by using Pearson correlation as distance for hierarchical agglomerative clustering with Ward’s method (“ward.D2”). Functional gene annotation was performed on each gene module individually using the Gene Ontology (GO_Biological_Process_2017) and the Kyoto Encyclopedia of Genes and Genomes (KEGG_2016) libraries with enrichR package^10^.

### 1.3. Mapping treatment-naïve ulcerative colitis and IBD risk genes to the murine RNA-seq dataset

To identify which genes are shared between mouse and human ulcerative colitis, we compared the list of DEGs identified by in the DSS dataset and the list of genes identified by Taman et al.^11^. Mapping of IBD risk genes was done using the list of IBD risk genes identified by fine-mapping at the single loci resolution ^12^. Identification of enriched GO processes and KEGG pathways was done using enrichR^10^.

### 1.4. Classification of ulcerative colitis molecular subtypes using genes in mouse principal components

To investigate whether the nuances of inflammation observed in the mouse model could also be found in humans, we made use of four human microarray datasets from GSE12251^13^, GSE73661^14^, GSE23597^15^ and GSE16879^16^. Combined, these datasets contain gene expression and metadata of 447 patients, containing information such as disease type (UC or CD), Mayo macroscopic score, the therapy given, when the sample was collected and the response to infliximab (IFX) or to vedolizumab (VDZ). Across all datasets, patients were considered inflamed if presenting a Mayo score of 2 or 3 (out of 3). Similarly, patient were considered to respond to therapy when it respective Mayo score reduced to 0 or 1, between 4-8 weeks of treatment with IFX or between 6-52 weeks of treatment with VDZ. For this study, we included only patients with UC before receiving any therapy (either IFX or VDZ), comprising a total transcriptional profiles of 143 patients, of which 102 received IFX and 41 for VDZ. The list of samples used in this study is supplied as metadata table (**Table S9**).

Probes with log2 fluorescence count lower than 6 in at least 10 samples were excluded from the analysis. Batches between dataset were observed and corrected using the ComBat function in SVA package^17^. Selection of genes for further exploration was done by different approaches: 1) using all genes; 2) using only the top 100 highly variable genes; 3) using the genes with top 100 high dispersion; 3) The gene with high loading in principal component 1 and; 4) The gene with high loading in principal component 2.

We determined whether clustering patterns exist by 4 independent methods: 1) By dimensionality reduction using tSNE. Since data originated from biopsies are known to present high variability across patients^18^, dimensionality reduction and visualization was done using t-Stochastic neighbor embedding (t-SNE). Because of it’s nonlinear characteristics, t-SNE becomes less sensitive to noise and outperform PCA^19^ to discriminate biopsies based on shared expression patterns, rather than their absolute expression values.; 2) By visual assessment of clustering tendency (VAT) using dissimilarity matrices^20^; 3) By using the Hartigan’s dip test^21,22^, which tests whether the gene distribution are different to an unimodal distribution. Values close to 1 indicate that the data is unlikely to present cluster substructures. We performed bootstrapping 100 times on 90% of the samples to calculate Hartigan’s dip test p-value. The comparison between bootstrapping with human highly variable genes and mouse PCs (see below) was done using paired Mann-Whitney test; 4) By dividing patients into subgroups using hierarchical agglomerative clustering. Cluster stability was determined by bootstrapping 300 times on 90% or the samples, resulting in the approximate unbiased (AU) statistics^23^. Clusters with AU closer to 100 present higher stability.

Instead of using the top variable genes as above, we alternatively used the top genes identified in the mouse RNA-seq DSS colitis dataset (see above). To this end, the top 100 genes identified in PC1 and PC2 were selected for identification of the respective human homologs. Together, 175 genes were found in top genes in both PC1 and PC2 and from these, 148 genes had a homolog in humans. In total, 57 homolog genes were found between our mouse PCs and the human dataset. Dimensionality reduction was performed with tSNE. Assessment of clustering tendency was done as described above. Agglomerative clustering on the Euclidean distance using complete linkage was used to discriminate patient subgroups UC1 and UC2. For the matter of definition used in this study, patients that present higher mean expression of the 57 mouse-human homologs were classified as UC1, while those with low expression were classified as UC2. Differences in expression between UC1 and UC2 were calculated using eBayes method in limma package^24^. Probes with fold changes above 1.5 and FDR lower than 0.001 were considered significantly differentially expressed. Identification of enriched GO, KEGG and cell types was done using enrichR^10^.

To identify which genes can discern UC1 from UC2, we trained a logistic regression classifier for each gene individually and comparing to the UC1 and UC2 classification mentioned above. The sensibility and sensitivity of the prediction was summarized using the area under the curve (AUC) method. Genes with AUC values closer to 1 (100%) have a better accuracy to distinguish UC1 and UC2 patients.

### 1.5. Lamina propria cell isolation for analysis by flow cytometry

Cell isolation from the colonic tissue was performed as previously described ^26^ with modifications. Briefly, tissues were open longitudinally, cut into 1cm pieces and washed with PBS. The epithelial cell fraction was obtained by incubating the tissue with Buffer-A (PBS, 5% FCS, 5 mM EDTA) at 37°C for 20 minutes under agitation at 600 rpm. The supernatant was collected and kept on ice while the remaining tissue was washed 2 times with PBS. Tissue were digested with collagenase solution containing 0.15 mg/ml Liberase TL (Roche) and 0.1 mg/ml DNase I (Roche) in HBSS and incubated at 37°C for 60 minutes under agitation at 1200 rpm. The digested and the epithelial cell fraction were mixed, filtered through a 100 um cell strainer, pelleted by centrifugation at 1750 rpm and re-suspended in Buffer-A. Cell suspensions were blocked with Fc-blocking solution (1:1000, eBioscience) and stained with the antibody mix (1:200), both at 4°for 15 minutes. The following antibodies were purchased from BD Biosciences: CD45.2 (104), CD3 (500A2), CD90.2 (53-2.1), EPCAM (G8.8), CD11b (M1/70), CD11c (N418), Ly6G (1A8), B220 (RA3-6B2) and CD64 (54-5/7.1). The following antibodies were purchased from eBiosciences: CD103 (2E7) and Ly6C (HK1.4). Counting beads (Spherotech) and DAPI (1:400, Sigma) were added to each sample to allow absolute cell quantification and exclusion of dead cells. Data acquisition was done using 5-laser LSR Fortessa flow cytometer (BD Biosciences) and analysis was carried out with FlowJo software (TreeStar).

### 1.6. Histological analyses

The colonic tissue was rinsed and flushed with PBS and gently squeezed out to remove non-adherent bacteria, fixed in 4% formaldehyde solution for 24 h and embedded in paraffin. 5 um sections were stained with H&E. Ki67 (1:100, Cat# MA5-14520, Thermo Scientific) staining was performed according to previously published protocol {26364605}. A pathologist accessed the tissue pathological score in a blind manner and score the sections as previously described^27^.

### 1.7. FITC-dextran assay

Assessment of epithelial barrier integrity was done as previously described. Mice were gavaged with 10 mg/mL FITC-dextran (Sigma) at different time points of DSS colitis, but on the same day of sacrifice. Four hours later, mice were killed and the blood collected for analysis. Sera were diluted 1:1 v/v in PBS and added to a 96-well plate for fluorescent-based assays (Invitrogen) and were quantified on a fluorescent plate reader using a 535/587nm ex/em filter. FITC-dextran concentration was calculated by interpolation to 12-dilution FITC-dextran standard curve.

### 1.8. Statistical analyses

Statistical analyses were performed using Prism Software 6.0 (GraphPad). Two-sample comparisons were compared using two-tailed Student’s *t*-test. ANOVA with Dunnett’s *post-hoc* was used for calculation of significance at multiple time points relative to the control (day 0). Non-continuous data was compared using non-parametric Mann-Whitney U test. Results were considered significant when p < 0.05.

### 1.9. Data availability

All the raw data generated in this study will be deposited in a suitable database (i.e., Gene Expression Omnibus) upon acceptance of this manuscript.

## Supplementary figure legends

**Figure S1.**
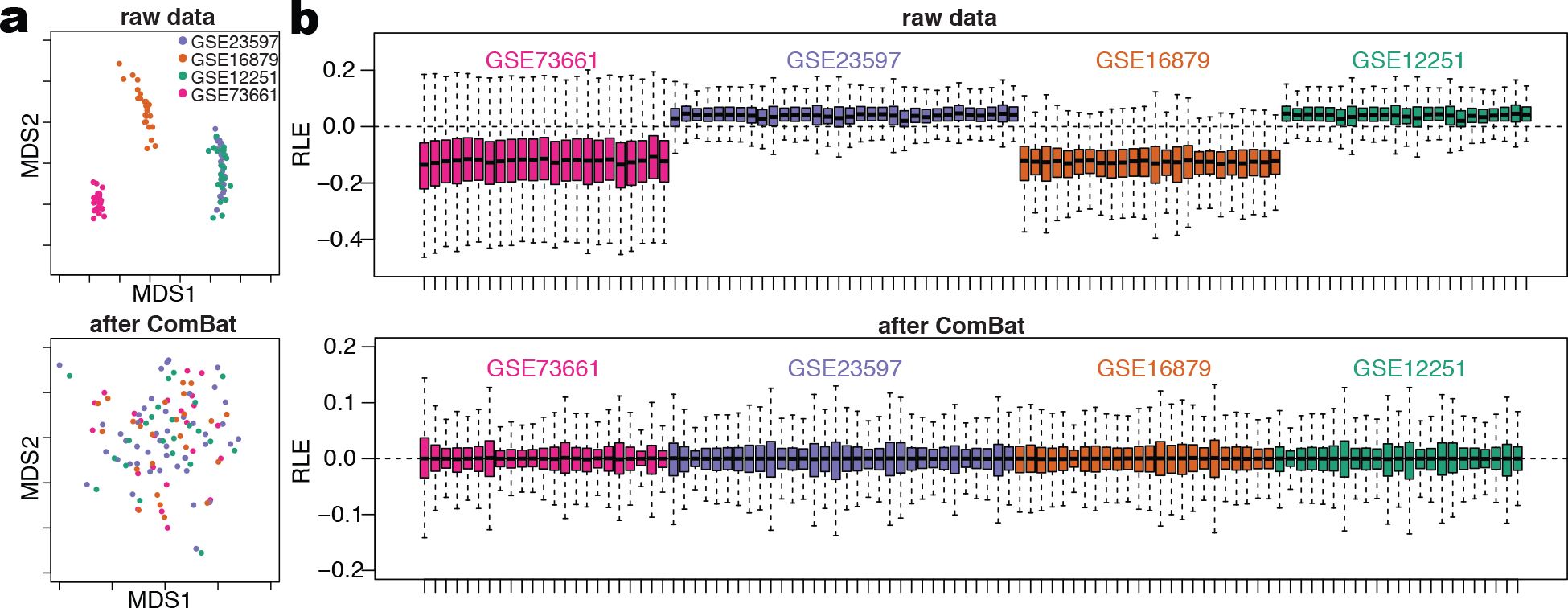
Normalization of publicly available ulcerative colitis datasets. (**a**) Multidimensional scaling plots before and after batch effect correction using ComBat. (**b**) Relative log expression plots comparing samples from the different datasets before and after adjusting for batches using ComBat.

**Figure S2.**
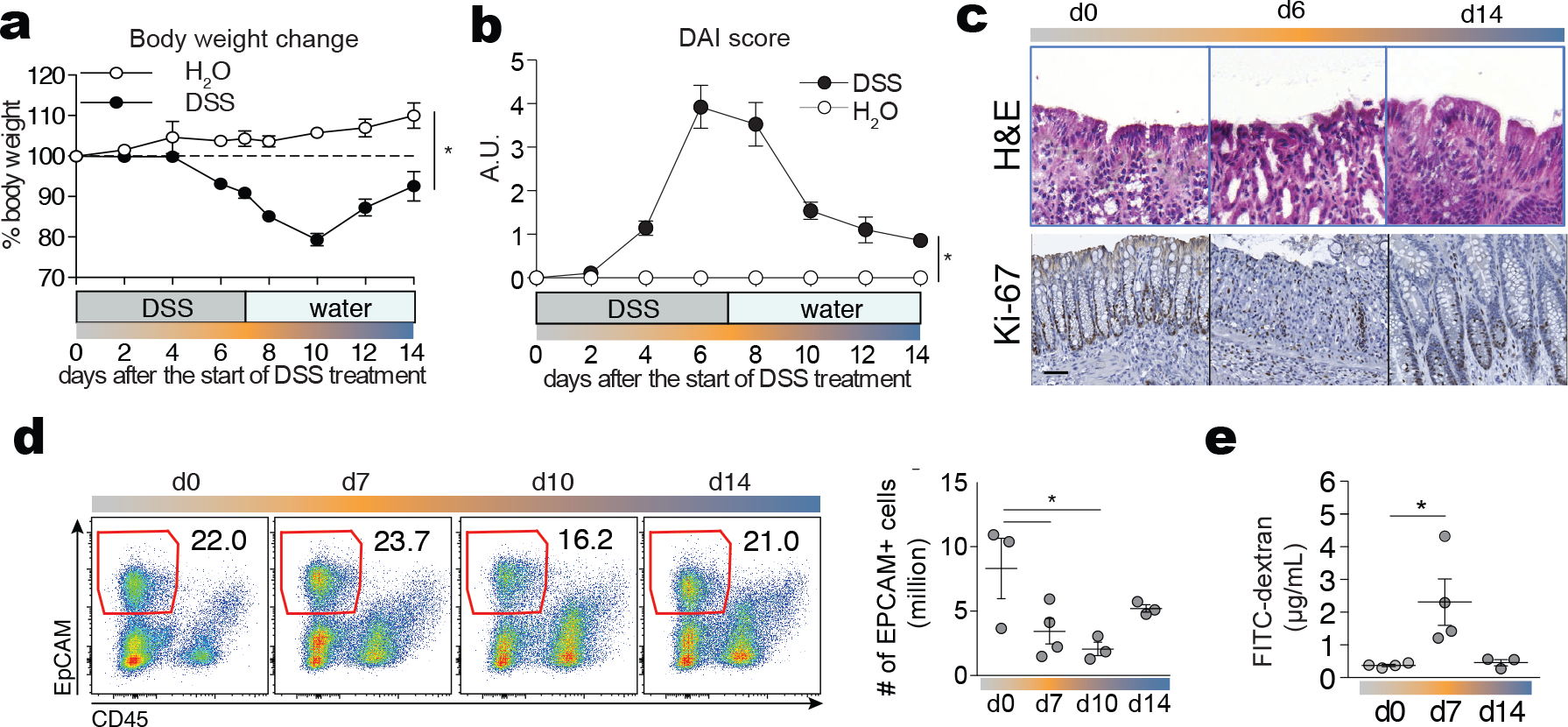
Macroscopic alterations in mice during DSS-induced colitis. (**a**) Body weight change over the time course of colitis. *P < 0.05; two-way ANOVA. (**b**) Disease activity index score (DAI) over time (in arbitrary units, A.U.). *P < 0.05; two-way ANOVA. (**c**) Representative histological section of the colonic tissue at indicated time points. H & E (upper) and immunohistochemistry staining for Ki-67 (bottom) are depicted. One representative figure out of three experiments. Scale bar 50 μm. (**d**) Flow cytometry data showing colonic epithelial cell (EpCAM^+^CD45^−^) frequencies during the course of the experiment. Dot plots are representative of three experiments. The graph on the right shows epithelial cell absolute numbers during the course of the experiment. *p < 0.05; two-way ANOVA. (**e**) Quantification of intestinal permeability by FITC-dextran assay. Mice were gavaged with 10 mg/mL of FITC-dextran and sacrificed 4 hours later for quantification of fluorescence in the serum. \**p* < 0.05; two-way ANOVA. Error bars represent SEM.

**Figure S3.**
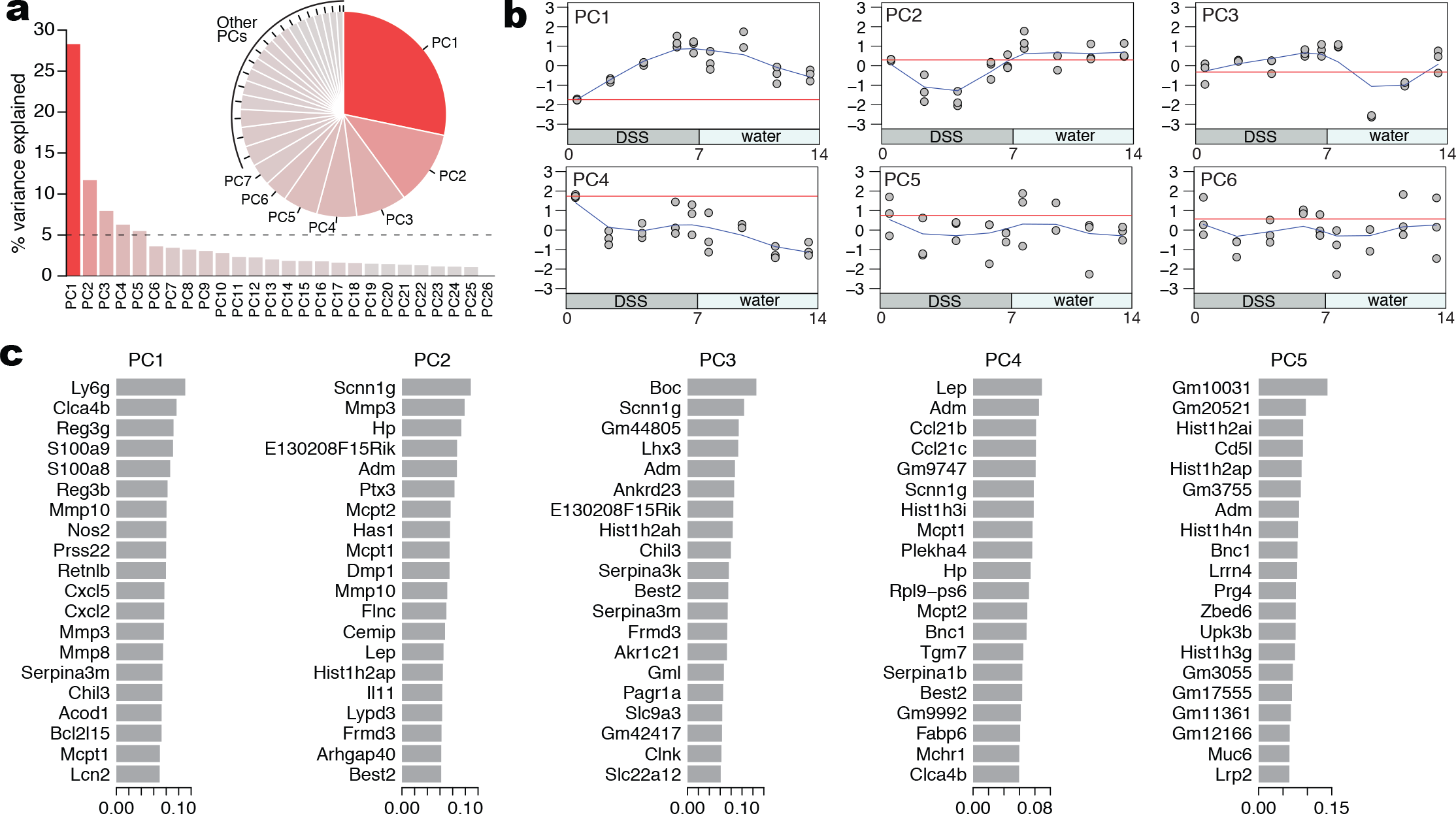
Identification of top leading genes and that drive overall differences in gene expression during DSS colitis. (**a**) Percentage of variance explained by each principal component (see Fig 2b). (**b**) Overall fluctuations in the first 6 PCs over the time course of DSS colitis. Note that the overall variance captured by PC6 is close to 0 and therefore not used in further analysis. (**c**) Ranking of the top 20 leading genes that contribute to the variance in each of the first 5 PCs.

**Figure S4.**
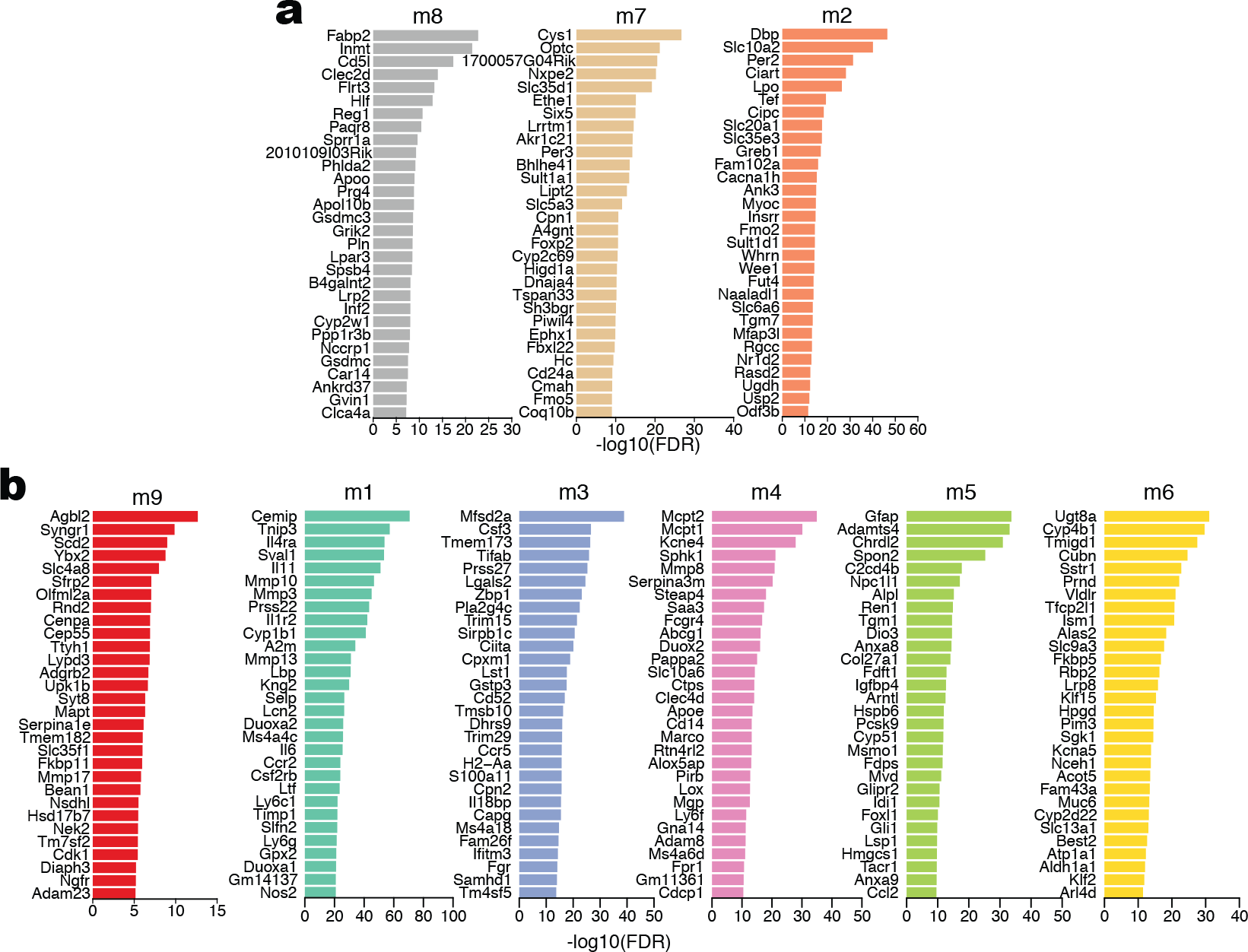
List of the top DEGs per module. (**a**) Down-regulated gene modules m8, m7 and m2 (see Fig 2c). (**b**) Up-regulated gene modules m9, m1, m3, m4, m5 and m6 (see Fig 2c).

**Figure S5.**
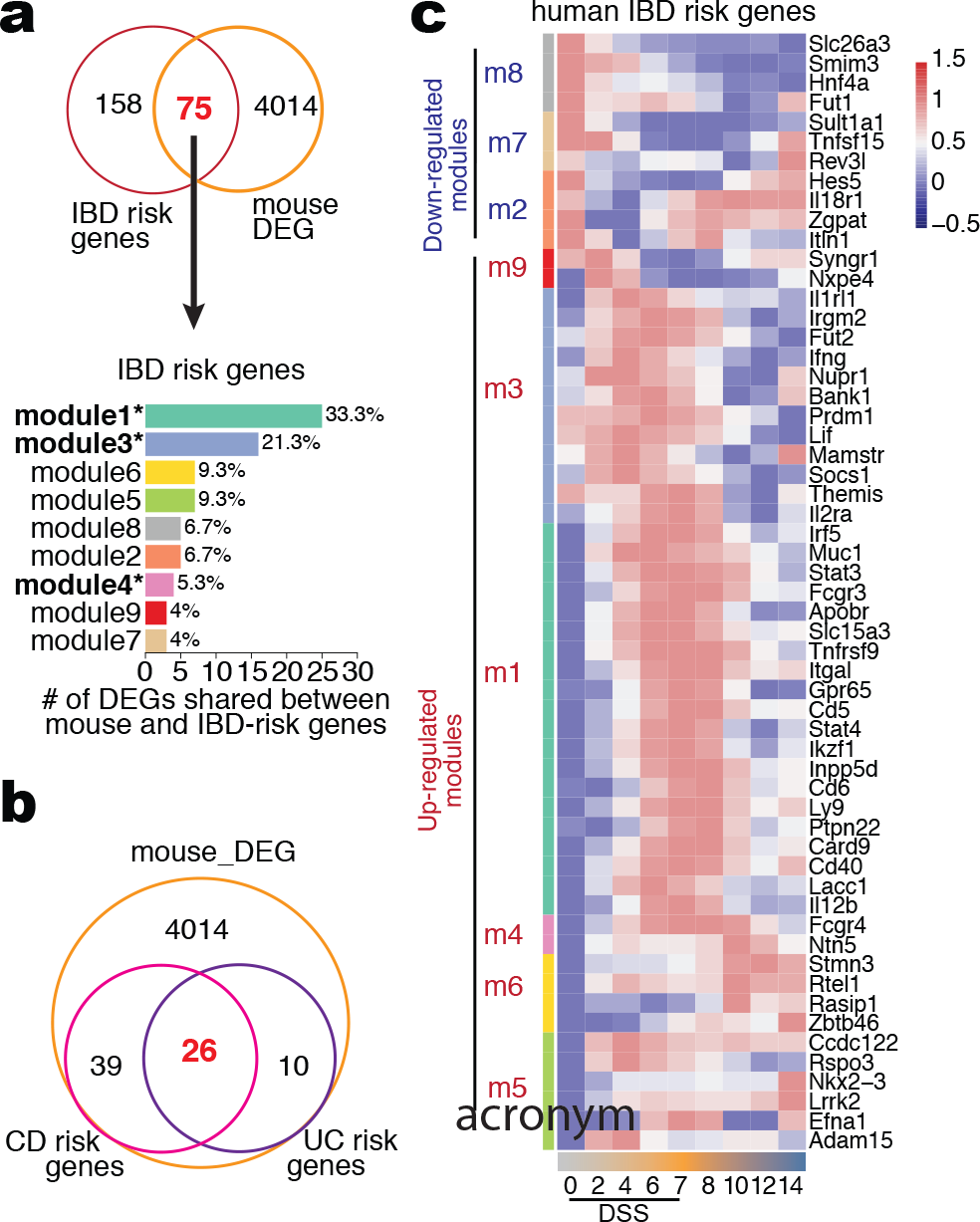
Mouse colitis and human UC share inflammatory pathways and IBD risk genes. (**a**) Venn diagram comparing IBD risk genes and the list of DEGs in the mouse dataset (upper). 75 genes are shared between these lists (in red). The number and percentage out of the 75 IBD-risk genes presented in each mouse module is shown (below). Modules highlighted in bold are the ones enriched for inflammatory terms in **Fig 2c**. (**b**) Venn diagram for the genes in the list of DEGs in the mouse dataset, genes associated with UC and/or to CD. Among those, 26 are shared between UC and CD (in red). (**c**) Expression level of IBD risk gene mouse homologs during the DSS colitis.

**Figure S6.**
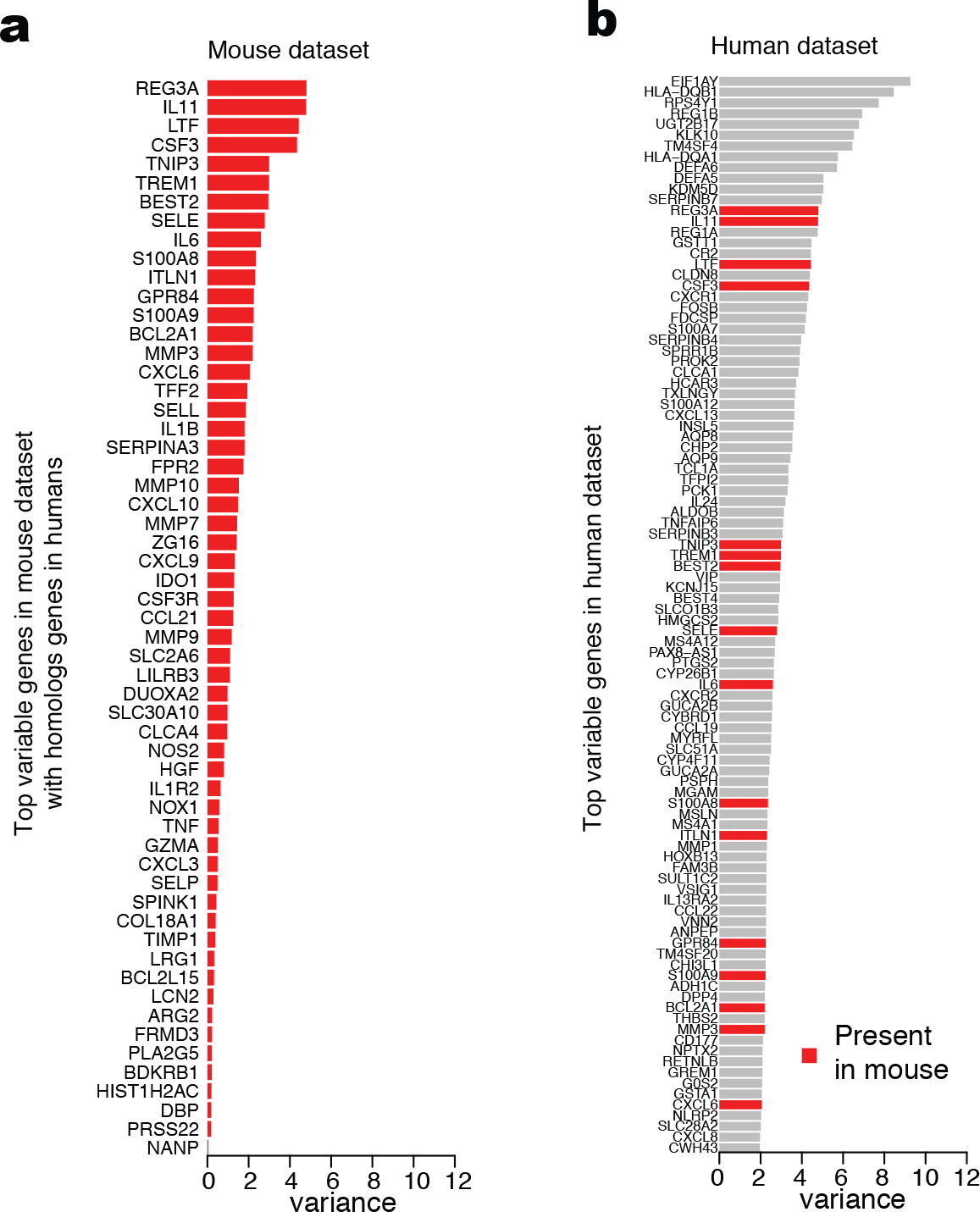
List of highly variable genes in humans and mouse colitis. (**a**) Top 100 genes sorted by high variance in the human dataset. Genes highlighted in red are also present among the top list of homolog genes identified in the mouse colitis dataset. (**b**) Top list of homolog genes identified in the mouse colitis dataset, sorted by variance on the human dataset.

**Figure S7.**
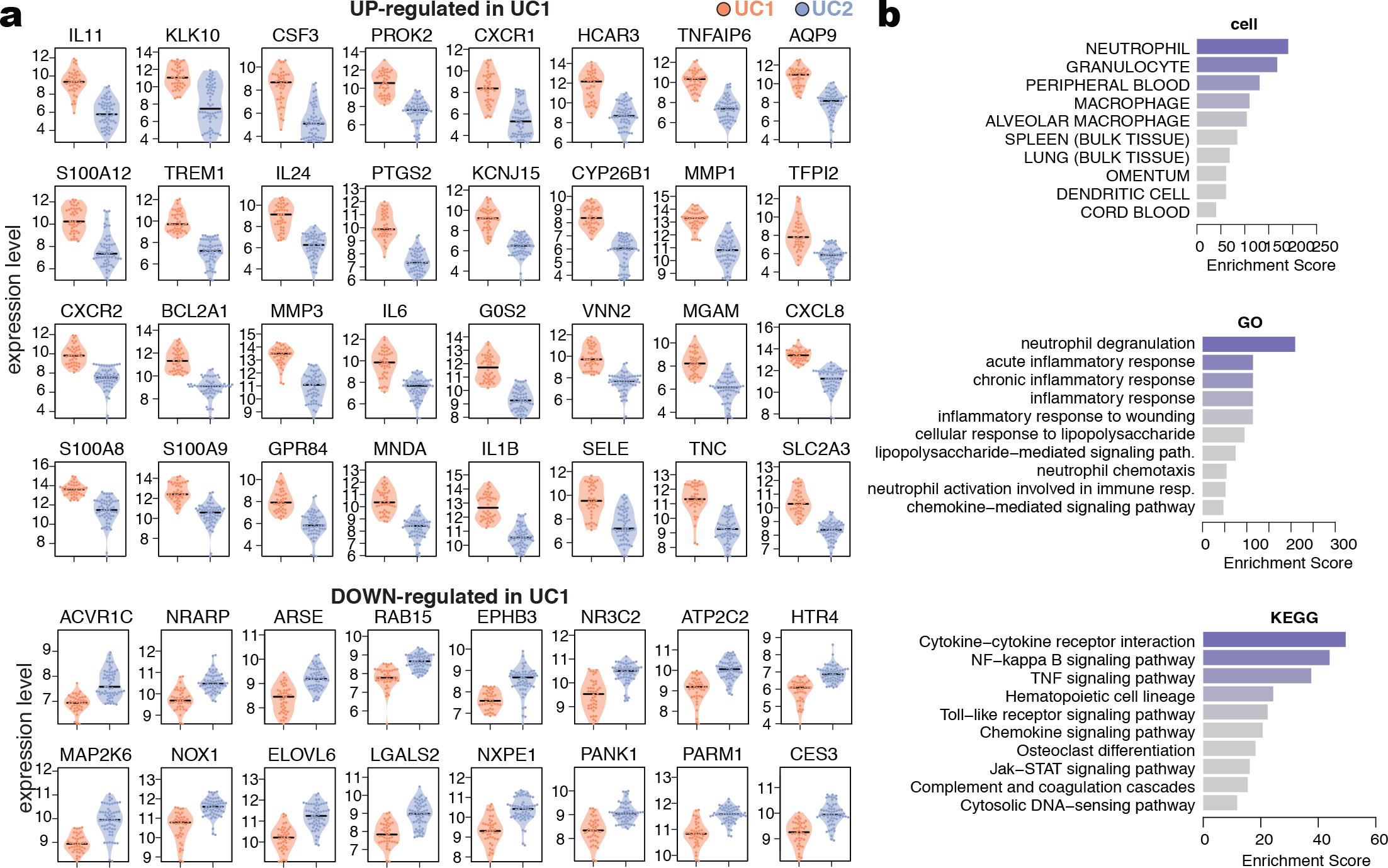
List of DEGs between UC1 and UC2 and enrichment analysis. (**a**) List of the top 32 up-regulated and 16 down-regulated DEGs between UC1 and UC2. (**b**) Cell, GO and KEGG enrichment analysis for the genes up-regulated in UC1 compared to UC2.

